# NeuroGPU: Accelerating multi-compartment, biophysically detailed neuron simulations on GPUs

**DOI:** 10.1101/727560

**Authors:** Roy Ben-Shalom, Nikhil S. Artherya, Alexander Ladd, Christopher Cross, Hersh Sanghevi, Kyung Geun Kim, Alon Korngreen, Kristofer E. Bouchard, Kevin J. Bender

## Abstract

The membrane potential of individual neurons depends on a large number of interacting biophysical processes operating on spatial-temporal scales spanning several orders of magnitude. The multi-scale nature of these processes dictates that accurate prediction of membrane potentials in specific neurons requires utilization of detailed simulations. Unfortunately, constraining parameters within biologically detailed neuron models can be difficult, leading to poor model fits. This obstacle can be overcome partially by numerical optimization or detailed exploration of parameter space. However, these processes, which currently rely on central processing unit (CPU) computation, often incur exponential increases in computing time for marginal improvements in model behavior. As a result, model quality is often compromised to accommodate compute resources. Here, we present a simulation environment, NeuroGPU, that takes advantage of the inherent parallelized structure of graphics processing unit (GPU) to accelerate neuronal simulation. NeuroGPU can simulate most of biologically detailed models 800x faster than traditional simulators when using multiple GPU cores, and even 10-200 times faster when implemented on relatively inexpensive GPU systems. We demonstrate the power of NeuoGPU through large-scale parameter exploration to reveal the response landscape of a neuron. Finally, we accelerate numerical optimization of biophysically detailed neuron models to achieve highly accurate fitting of models to simulation and experimental data. Thus, NeuroGPU enables the rapid simulation of multi-compartment, biophysically detailed neuron models on commonly used computing systems accessible by many scientists.

## Introduction

Electrical activity of single neurons is determined by the distribution of various ionic conductances arranged across complex morphologies (Mainen and Sejnowski, 1996a; Hausser et al., 2000; London and Häusser, 2005; Spruston, 2008; Hay et al., 2013; Alonso and Marder, 2019). Our understanding of single neurons has long relied on the ability to construct biophysically rigorous models of how neuronal membrane potential [Vm], and, hence, action potentials (APs, spikes) are generated from currents [I] flowing across the membrane and through the cell (Fig. 1) (Hodgkin and Huxley, 1952). Wilfrid Rall described the biophysical theory of how membrane potential of a single neuronal segment (‘compartment’) depends on the conductance (e.g., gNa) and voltage dependent flow of specific ionic species [e.g., sodium (Na) and potassium (K)], as well as passive properties of the membrane (i.e., capacitance) (Fig.1, top row) (Rall, 1962a). Using cable theory, Rall further described how to connect different compartments of a neuron, providing the foundation for modeling complex, spatially extended neuronal morphologies (Fig.1A) (Rall, 1962b). Concomitantly, the membrane channels that mediate a specific ionic current exhibits large diversity of genetically defined conductances (e.g., gNav1.2, gNav1.6, etc.,), indicating that individual compartments are, in reality, quite complex (Fig.1B). While just beginning to be appreciated at the time, we now know that ion channels have their own voltage dependent kinetics that are determined by the transition probabilities amongst various states of the channel subunits (Fig.1C) (Hille, 1984; Colquhoun and Hawkes, 1995). Finally, simulations of realistic neural network models require faithfully capturing the complexity of the individual neurons (Einevoll, et al., 2019). While the physical theories required to link these vastly disparate spatial-temporal scales exists, our ability to utilize them for basic understanding and clinical translation is impeded by the computational burden of the required simulations (Fig.1D, E).

**Figure 1:**
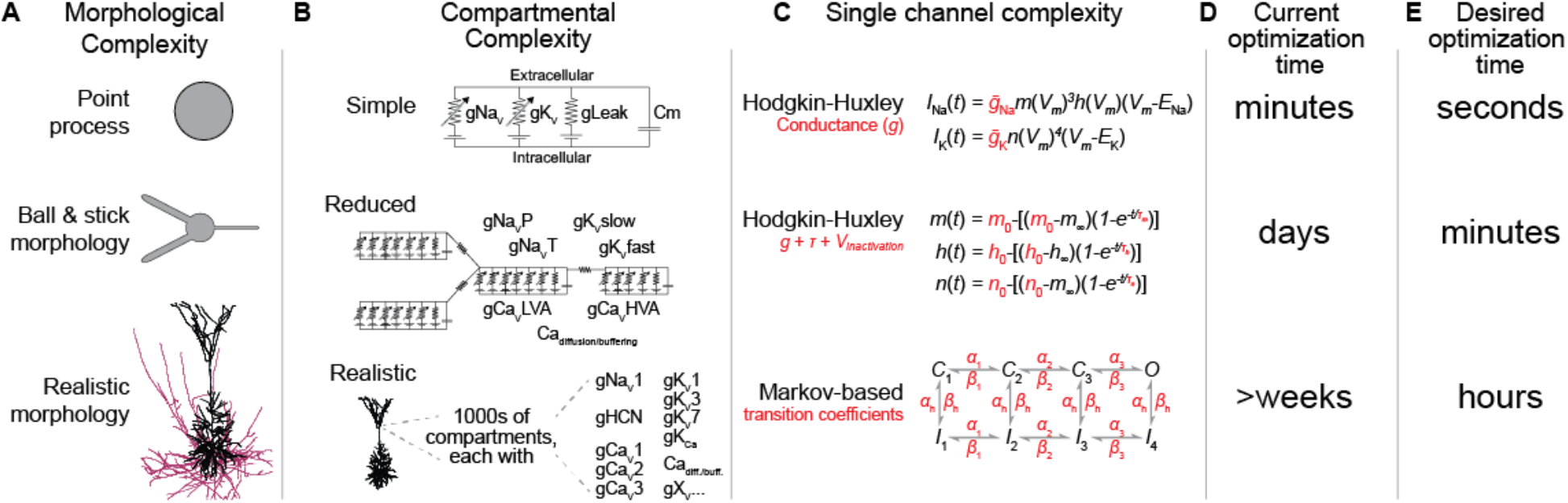
Optimizing biophysical neuronal models increase with their complexity. Biophysical neuronal model complexity depends on three factors (A-C) which determine computational costs required to simulate and fit models to empirical data (optimization) (D-E) **A:** Morphological complexity: The abstraction of the neuronal morphology ranges from a point neuron, to a simplified morphology, and ultimately toward different methods to reconstruct neuronal morphology with high resolution. **B:** Compartmental complexity: Compartmental models vary in complexity both in number of compartments and the content of each compartment. Top: Single compartment with basic, conductances required for spiking and a resting membrane potential. Middle: Multi-compartmental model where ion-channels are aggregated under several conductances e.g. all different voltage gated potassium channels are represented in a slow and fast inactivating channel (Korngreen and Sakmann, 2000). Bottom: High resolution morphology with detailed representation to all channels sub-types in each compartment. **C:** Single-channel complexity: The overall conductance in compartmental models is formalized either with Hodgkin-Huxley equations or with Markov-based models. When fitting a model to empirical data, several parameters can be varied. Top: only the maximal conductance in a Hodgkin-Huxley formulation. Middle: coefficients added to Hodgkin-Huxley equations (time constants, voltage dependence). Bottom: In Markov-based channels the transition coefficients can be varied. **D:** Optimizing a model (fitting model to data) depends on the complexity of the model and number of free parameters. When complexity and number of free parameters require compute times that exceed a few days, it becomes impractical to use high resolution models. **E:** Developing tools to accelerate simulation and optimization time will enable us to use more complicated and biophysical relevant models.

Our predictive understanding of several neuronal cell classes has benefitted from the repeated refinement of compartmental models that describe their activity in health and disease. For example, neocortical pyramidal cells have been modeled extensively, providing insight into the effects of morphology on AP firing characteristics (Mainen and Sejnowski, 1996b; Hay et al., 2011; Almog and Korngreen, 2014), how synaptic input patterns affect spike output (Maršálek et al., 1997; Dlesmann et al., 1999; Destexhe et al., 2003) how APs initiate and propagate within axons (Kole et al., 2007, 2008; Shu et al., 2007; Hu et al., 2009; Hallermann et al., 2012; Cohen et al., 2020), and how neuronal activity is affected by alterations in ion channel function induced by genetic variation (Zamponi et al., 2010; Allen et al., 2014; Ben-Shalom et al., 2017; Spratt et al., 2019). Similar intensive studies have focused on other cell classes, including hippocampal pyramidal cells (Mainen et al., 1996; Magee and Cook, 2000; Poirazi et al., 2003; Narayanan and Johnston, 2008; Milstein et al., 2015)), cerebellar Purkinje cells (De Schutter and Bower, 1994; Miyasho et al., 2001; Roth and Häusser, 2001), and midbrain dopaminergic neurons (Canavier, 1999; Canavier and Landry, 2006; Kuznetsova et al., 2010). In parallel with these advances in modeling, there has recently been an enormous improvement in experimental approaches to better understand the diversity of neuronal classes and their activity patterns. Within the general group of neocortical pyramidal cells, for example, exists a wealth of diversity. This includes not only differences in morphology and activity across laminae (Smith and Häusser, 2010; Deitcher et al., 2017; Kanari et al., 2019), but also within laminae depending on genetic makeup or axonal projection targets (Dembrow et al., 2010; Gee et al., 2012; Clarkson et al., 2017), or even within what was thought to be a homogenous cell class within a single layer as one samples across a large region of cortex (Fletcher and Williams, 2019).

Given the enormous complexity and vast spatio-temporal scales described above, generating models that accurately recapitulate neuronal activity across the true range of diversity present in nature can be a daunting task. Model fitting often requires one to tune individual parameters to best match empirical observations. This process can be aided by iterative rounds of parameter exploration and optimization that aim to minimize the differences between empirical data targets and their associated models. These procedures can be computationally demanding (Fig.1D-E). Indeed, each linear improvement in model accuracy requires an exponential increase in computational resources (Nocedal and Wright, 2006; Gurkiewicz and Korngreen, 2007). Thus, model optimization is often done on supercomputers that parallelize these computations across massive number of central processing unit (CPU) cores. Unfortunately, the cost of constructing and operating supercomputing centers is similarly massive. As such utilization of these resources are typically restricted to large consortia, such as the Blue Brain Project (BBP) (Markram et al., 2015) and the Allen Institute (Gouwens et al., 2018). For more restricted budgets, simulations must typically be compromised in scale, complexity, or accuracy, thus negatively impacting results (Almog and Korngreen, 2016).

In the past 10 years, graphics processing units (GPUs) have emerged as an alternative to CPU-based hardware that may offer comparable levels of performance at substantially reduced cost for some problems. GPUs utilize streaming multiprocessors with multiple simple cores that allow for distributed, parallelized computing for relatively small chunks of data. With software optimized GPUs, GPU-based computing can often outperform CPU-based applications in processing speed and cost for some problems (Payne et al., 2010). Today, GPUs are being used in scientific simulations, including molecular dynamics (Go et al., 2012; Salomon-Ferrer et al., 2013) and climate modeling (Prein et al., 2015), and are the computational engine for most modern artificial intelligence applications (Schmidhuber, 2015). In neuroscience, GPUs are currently being used to accelerate complex imaging dataset processing (Eklund et al., 2013), spiking neural network analysis (Fidjeland and Shanahan, 2010), clustering of activity from *in vivo* extracellular electrophysiological experiments (Pachitariu et al., 2016), and simulations of single ion channels (Ben-Shalom et al., 2012). Recently two platforms for neuronal biophysical simulations with GPU support were developed: Arbor (Akar et al., 2019) and CoreNeuron (Kumbhar et al., 2019). Both platforms focused on simulating large scale neuronal networks comprised of detailed multi-compartmental models. Each approach has unique advantages and disadvantages. For example, CoreNeuron supports previous NEURON models, but is not implemented in CUDA, the fundamental operating language of NVIDIA’s GPUs. As such, its ability to accelerate model runtimes with GPUs is relatively modest. Arbor, instead, is implemented in CUDA via an entirely new simulation environment. Thus, while it does harness the speedup potential of GPUs, it is not clear how existing models, such as those found in ModelDB and the BBP portal (McDougal et al., 2015; Ramaswamy et al., 2015), could be ported to Arbor, thus impeding utilization.

Here, we describe NeuroGPU, a computational platform optimized to exploit GPU architecture to dramatically accelerate simulation of mutli-compartmental neuronal models. Our goal with NeuroGPU was different than that of CoreNeuron or Arbor. Rather than focusing on neuronal network simulations, NeuroGPU is designed to optimize fitting of models that best recapitulate empirical data derived from single neurons. To do so, we developed new approaches to parallelize compartmental models, utilizing the GPU-based programming language CUDA to optimize memory handling on GPUs. This resulted in simulation speedups of up to 200-fold on a single GPU and up to 800-fold using a set of 4 GPUs. Building on our previous efforts (Ben-Shalom et al., 2013), we developed an intuitive user interface that can import most compartmental models implemented in NEURON (Carnevale and Hines, 2006) deposited at the ModelDB (McDougal et al., 2017) or BBP portal (Ramaswamy et al., 2015). Further, we provide methods to explore model parameter space to study how each parameter of the model contributes to its voltage output. The runtimes of NeuroGPU enables us to sample the parameter space in a very detailed manner, which were utilized here to reveal the response landscape of single neurons. Finally, we provide an interface to use NeuroGPU for fitting models to experimental data with evolutionary algorithms (DEAP and BluePyOpt) (Gagn, 2012; Van Geit et al., 2016). NeuroGPU implemented such model optimization algorithms in minutes, rather than days, which had been the previous benchmark (Hay et al., 2011; Almog and Korngreen, 2014). This enables the use of more complex models that will better represent experimental data, as we demonstrate here in both simulated and experimental data. NeuroGPU therefore provides an open-source platform for neuronal simulation with increased speed and reduced cost, thus enabling the neuroscience community to perform high quality biophysically detailed simulations.

## Methods

### Hardware

NEURON and TitanXP-based simulations were run on a PC with Intel Core I7-7700K 4.2GHz with 16GB of RAM. Tesla V100-based simulations were run using the NVIDIA PSG cluster. Here, each simulation was run on a single node with Haswell or Skylake CPU cores. For multi-GPU simulations, we used cluster nodes with NVLINK (Li et al., 2019) between the GPUs to enable memory peer-access. Fitting models to experimental data was done on the Cori GPU cluster from the National Energy Research Scientific Computing Cent (NERSC) at Lawrence Berkeley National Labs. Cori GPU nodes includes 8 NVIDIA Tesla V100 and 20 Skylake CPU cores with total of 384GB memory.

### Software

Simulations were performed in NEURON 7.6 and CUDA 10.1. All scripts were written in Python 3.7. All software is available at https://github.com/roybens/NeuroGPU.

### Importing NEURON models

The python script extractmodel.py exports NEURON models to NeuroGPU. This script reads all simulation details from runModel.hoc, which is populated using the GUI (Fig. S1). NEURON models are described using either hoc or python scripts. The scripts include a morphology that can either be called as a separate file or constructed within the script (Fig S1B). The user must input a file containing model stimulation, which includes temporal aspects of the model and command currents delivered at a prescribed location. Furthermore, all free parameters, such as channel properties, must be described. These import components are translated into CUDA code, termed kernels, that can run on the GPU via the python script “extractmodel.py” (Fig S1C). This script first takes runModel.hoc and loads it into NEURON, not to run simulations, but rather to query NEURON for model properties needed for subsequent porting to NeuroGPU, including compartment names and the tri-diagonal matrix (F-Matrix or Hines Matrix) which holds the differential system for the voltages of the dendritic tree. Then, the script iterates over the .mod files in the directory, parses them and creates relevant kernels for each mechanism described. Mechanism kernels are written to the AllModels.cu in similar structure as described previously (Hines and Carnevale, 2000; Carnevale and Hines, 2006), iterating over all compartments defined in the model. A new hoc file is created to register mechanism values, which are stored in AllParams.csv and inserted in each compartment. After model parameter maps are determined, they are cataloged as part of ParamMappings.txt for reference for future reiterations of the same model, eliminating the need to reload NEURON. Finally, the script writes code translated to CUDA in NeuroGPU.cu and packages the application to run on either Windows or Unix. After compiling the code, an executable is created that reads the AllParams.csv and the stimulation and runs the model on the GPU.

### Translating mechanisms to CUDA and memory assignment

Mechanisms in NEURON are described by NMODL (.mod) files (Hines and Carnevale, 2000), which update the mechanism states every simulation time step. This is done using three different procedures within NEURON that initialize mechanisms (nrn_init), update currents that mechanisms affect (nrn_cur), and then update mechanism states (nrn_state) (Carnevale and Hines, 2006). In NeuroGPU, CUDA kernels are written for each of these procedures using .mod and .c files that are generated by NEURON when running nrnivmodl. Kernels are saved and editable in AllModels.cu and AllModels.h.

CUDA is an extension of the C programming language that enables computation on the GPU (Nvidia, 2018). CUDA kernels are procedures running on the GPU that can be invoked from either the GPU or CPU. To invoke a kernel from the CPU, one must specify the number of parallel threads used. Threads, which allow for parallelization on the GPU, are organized into blocks, with each thread occupying a specific address within that block (idx.x) (Fig S2 C). GPUs are structured to operate well when computing 32 parallel threads, a computing structure termed a warp (Nvidia, 2018). Therefore, we structured NeuroGPU to utilize 32 threads in the x dimension, corresponding to individual morphological segments within the model. For a given model with more than 32 segments, individual threads are responsible for calculating every 32^nd^ segment. For example, thread #1 would calculate segments 1, 33, 65, ... 31N+1.

Complex neuronal models, including many described in the BBP portal, are memory intensive (Hay et al., 2013; Ramaswamy et al., 2015). GPUs have several forms of memory that have tradeoffs in terms of their size and relative speed that make them ideal for certain aspects of model processing and impractical for others. GLOBAL memory is the largest physical memory space available on the GPU but is relatively slow. Here, we use GLOBAL memory to store the largest data structures associated with a given model, in part because they simply cannot be held by other memory structures. SHARED memory is far faster, shared among the whole GPU block, but limited to 48 kilobytes. This makes it ideal for storing the tridiagonal matrix, as this matrix is the most accessed data structure within NeuroGPU. CONSTANT memory, which is a 64 kilobyte block of fast, read-only memory, is used to hold constant data structures, including the order in which the tri-diagonal matrix is solved in parallel (Ben-Shalom et al., 2013). Lastly, REGISTER\LOCAL memory is the fastest memory available on the GPU but is limited to maximum of 63 registers per thread and a total of 16 kilobytes of memory shared across the entire block. It is used to store local variables necessary for the course of the simulation (Fig. S2C).

### Extracting simulation properties from NEURON

NeuroGPU utilizes NEURON for simulation pre-processing, including mechanism translation (which includes mathematical descriptions of various ion channels, calcium diffusion characteristics, and other elements of neuronal function, (Hines and Carnevale, 2000), a map for mechanism distribution across compartments (ParamMappings.txt), and exporting the tri-diagonal matrix using fmatrix(). These are stored in BasicConstSegP.csv. NEURON extracts all parameters for cable equations and mechanism values within each compartment to AllParams.csv (Fig. S1). External stimulation delivery location, intensity, and timecourse are written in stim.csv. Resting membrane potential and number of time steps in the simulation are written in sim.csv.

### Solving the tridiagonal matrix

Matrix solutions were performed using the branch-based parallelism approach as described previously (Ben-Shalom et al., 2013), with morphology analysis guiding iterative matrix computations. This analysis is done in extractmodel.py and the data structures to solve the tri-diagonal in parallel is stored in BasicConstSegP.csv.

### Benchmarking

All benchmarking was done compared to NEURON 7.6 running in a single thread. The morphology was adjusted to have one segment per compartment in both NEURON and NeuroGPU comparison. Simulation runtimes were compared without hard drive read/write file steps, as these aspects depend more on hard drive properties than CPU/GPU comparisons.

### Multi-compartmental models

NeuroGPU performance was tested with 4 different models:

1. A passive model, utilizing passive channels described in NEURON distribution pas.mod file. These channels were distributed on both simple and complex morphologies (see Fig. S3A, D) (Mainen and Sejnowski, 1996b). The simple morphology was based on the simple morphology described in Mainen and Sejnowski, with compartments reduced to 32, as this is the minimum number of compartments required for NeuroGPU-based simulations.
2. The Mainen and Sejnowski (1996) model, with channels distributed on the same complex and simple morphologies. Channels are distributed as in (Mainen and Sejnowski, 1996b) (Fig. S4)
3. A pyramidal cell model from the Blue Brain Project portal (Ramaswamy et al., 2015) (Fig. 2). BBP_PC refers to the model named L5_TTPC1_cADpyr232_1.
4. A chandelier cell model, termed BBP_CC, referring to L5_ChC_dNAC222_1. For this model, the Kdshu2007.mod files were altered to run on NeuroGPU. Specifically, global variables were removed from the neuron block and instead placed in the assigned block (Carnevale and Hines, 2006) (Fig. 2).

**Figure 2:**
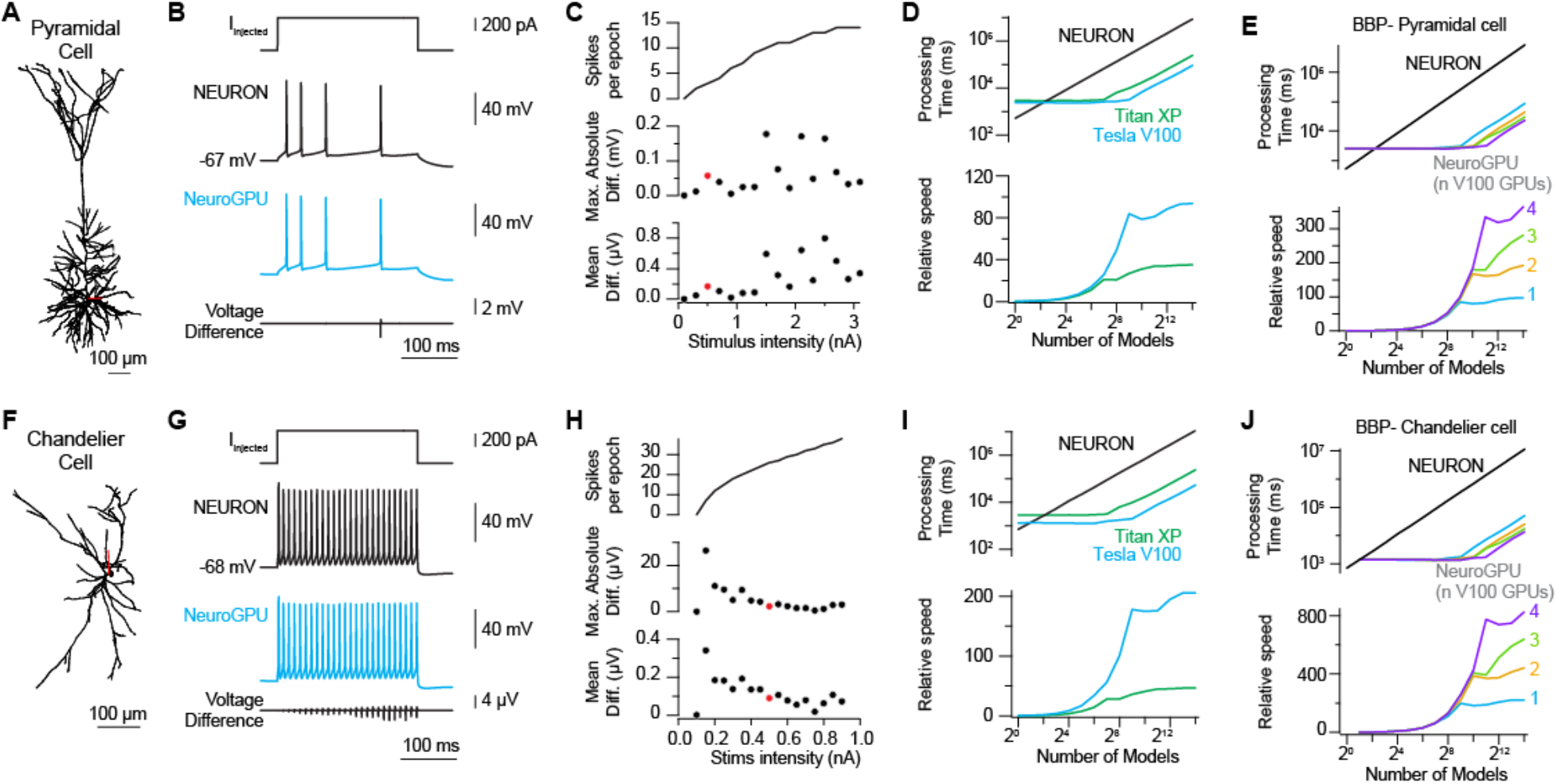
NeuroGPU reduces simulation run-time of complex neurons by orders of magnitude without compromising accuracy. **A:** Morphology of a BBP portal layer 5 neocortical pyramidal cell (Ramaswamy et al., 2015). Dendrite in black, axon in red. **B:** Top: injected current at the soma. Middle: NEURON voltage response as recorded at the soma. Cyan: NeuroGPU response as recorded at the soma. Bottom: difference in voltage between NEURON and NeuroGPU. **C:** Top: APs generated per current injection intensity in the soma. Middle, bottom: Peak and average voltage difference between the voltage response in NEURON and NeuroGPU. Red circles denote examples in B. **D:** Top: Runtimes for the model using the different architectures: black – NEURON, green – NeuroGPU on TitanXP, blue – NeuroGPU on TeslaV100. X-axis in log2 scale, Y-axis in log10 scale. Bottom: Speedup compared to NEURON. **E:** Top: Runtime for the model on 1-4 GPUs (Tesla V100) on the same node. Bottom: Speedup compared to NEURON **F-J:** Same as A-D, but for a model chandelier cell.

### Optimization algorithms

Two different genetic algorithm versions were used in this study. For data related to Figure 4, the *eaMuPlusLambda* algorithm from the DEAP package was implemented by modifying the varOR procedure to call NeuroGPU (Rainville et al., 2012). Optimization was performed on the BBP_PC model. For each iteration, the algorithm began with a new population of parameters with values randomly chosen with the range specified in Table 1. The model was modified to accept new values from the optimization algorithm (similar changes were necessary for parameter space exploration for Figure 3). Target data were generated using the original parameters values described in Table 1. Optimization was targeted to reduce error between target data and test data using both the interspike interval (ISI) and the root mean square (RMS) of the voltage as the error function. Error was reduced to a single variable by weighting these two variables as: 10*ISI + RMS.

**Table 1.** – parameters varied in the BBP model for figure 4:

| Parameter Name | Base value | Lower Bound | Upper Bound |
| --- | --- | --- | --- |
| gNaTa_tbar_NaTa_t | 3.137968 | 0.3137968 | 31.37968 |
| gNaTs2_tbar_NaTs2_t | 0.983955 | 0.0983955 | 9.83955 |
| gK_Tstbar_K_Tst | 0.089259 | 0.0089259 | 0.89259 |
| gIhbar_Ih | 0.00008 | 0.000008 | 0.0008 |
| gImbar_Im | 0.000143 | 0.0000143 | 0.00143 |
| gSKv3_1bar_SKv3_1 | 0.303472 | 0.0303472 | 3.03472 |

**Figure 3:**
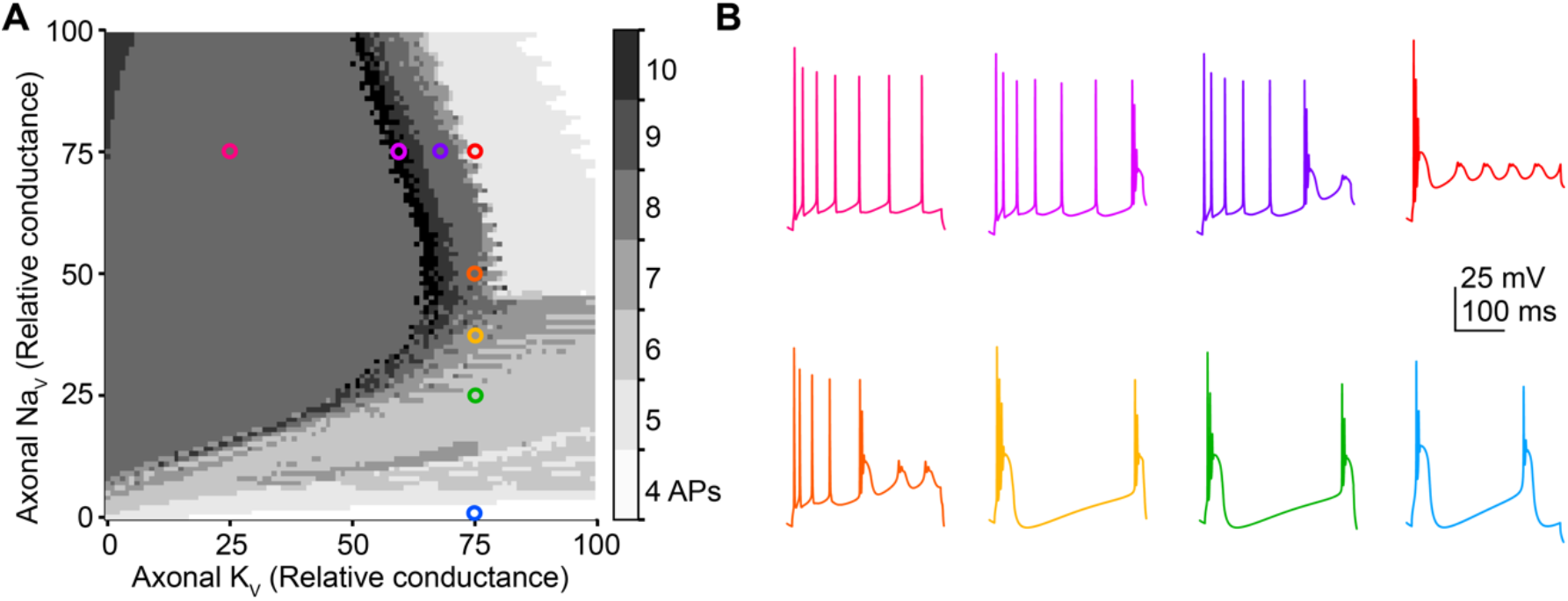
NeuroGPU enables rapid exploration of *parameter space in complex pyramidal neuron model*. **A:** Each point in the grid represents the number of APs in the relevant model. Points on the axis represent the varied conductances of Na_v_ and K_v_ at the axon in the range of [0,10] and [0,20] S/cm^2^, respectively. **B:** Example voltage responses for chosen models from A. Colors match to the corresponding model location.

For data related to Figure 5, the BluePyOpt (Van Geit et al., 2016) implementation of Multiple Objective Optimization (MOO) was used. Experimental target data for these experiments were from whole-cell current-clamp recordings from layer 5b thick tufted pyramidal cells in acute slices from wild-type mouse prefrontal cortex (postnatal day 62) (Spratt et al., 2019). Optimization was targeted to minimize the root mean square voltage error at each time point between empirical data and model output as well as the following objectives, as defined in the electrophysiology feature extraction library (eFEL) from the BBP (Moor et al., 2015): voltage_base, AP_amplitude, voltage_after_stim, ISI_values, spike_half_width, and afterhyperpolarization_depth. Electrophysiological data were fitted in models with morphology from L5_TTPC1_cADpyr232 Fig (5A) or a reconstructed prefrontal cortex L5 thick tufted pyramidal neuron deposited at neuroMorpho.org (Ascoli et al., 2007; Yin et al., 2018). The model parameters that were varied for the S1 model and PFC model are described in Table 2 & 3 respectively.

**Table 2.** parameters varied in the BBP model for figure 5A:

| Parameter Name | Base value | Lower Bound | Upper Bound |
| --- | --- | --- | --- |
| g_pas_all | 0.0003 | 0.000003 | 0.003 |
| e_pas_all | -75 | -150 | -50 |
| gNaTa_tbar_NaTa_t_axonal | 3.137968 | 0.3137968 | 31.37968 |
| gK_Tstbar_K_Tst_axonal | 0.089259 | 0.0089259 | 2.89259 |
| gNap_Et2bar_Nap_Et2_axonal | 0.006827 | 0.0006827 | 0.06827 |
| gK_Pstbar_K_Pst_axonal | 0.973538 | 0.0973538 | 9.73538 |
| gSKv3_1bar_SKv3_1_axonal | 1.021945 | 0.1021945 | 10.21945 |
| gCa_LVAstbar_Ca_LVAst_axonal | 0.008752 | 0.0008752 | 0.08752 |
| gCa_HVAbar_Ca_HVA_axonal | 0.00099 | 0.000099 | 0.0099 |
| gSKv3_1bar_SKv3_1_somatic | 0.303472 | 0.0303472 | 3.03472 |
| gCa_HVAbar_Ca_HVA_somatic | 0.000994 | 0.0000994 | 0.00994 |
| gNaTs2_tbar_NaTs2_t_somatic | 0.983955 | 0.0983955 | 9.83955 |
| gCa_LVAstbar_Ca_LVAst_somatic | 0.000333 | 0.0000333 | 0.0333 |
| gIh_dend | 0.00008 | 0.000007 | 0.009 |

**Table 3.**
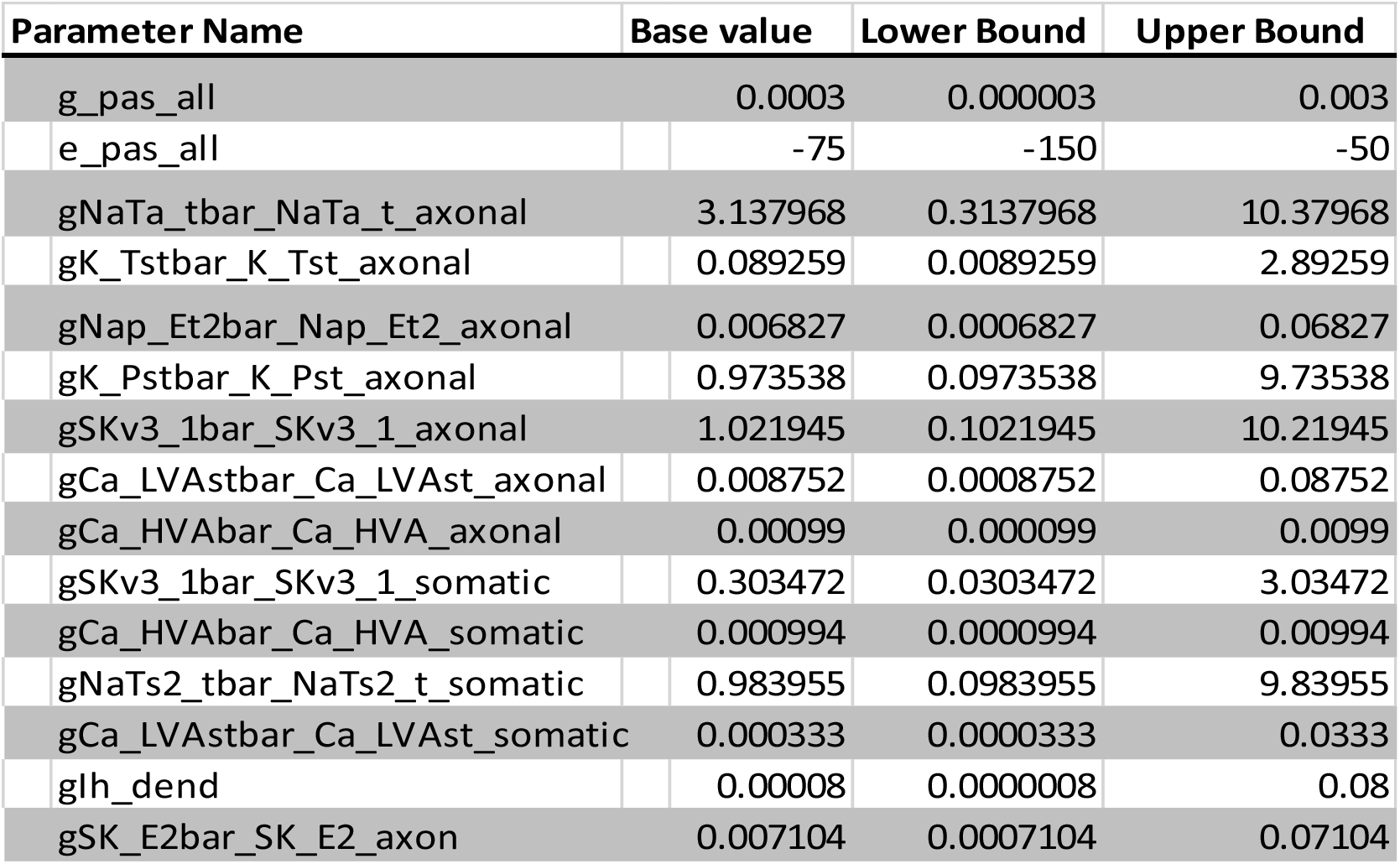
parameters varied in the BBP model for figure 5B:

### Profiling

To better understand why NeuroGPU accelerated some models more than others, we used the NVIDIA profiler to monitor GPU utilization. Further, we tested two different memory handling configurations to determine how best to utilize GPU parallel processing. In both cases, the GPU is responsible for updating ionic currents from established mechanisms, solving the tridiagonal matrix, and updating model states and voltages at each time step. In the first configuration, termed SingleKernel, we computed all the time steps of each simulation in one kernel on the GPU, largely because this would limit the amount of time performing the relatively slow step of transferring memory between the GPU and CPU. In this case, the transfer is done only once and during the simulation the GPU communicates with the CPU only to transfer voltages reported at the recording electrode site. Alternatively, we also created a SplitKernel condition, in which the simulation is split into many small kernels that are invoked every single time step. Data are then registered back to the CPU and the next time step is run in serial. This approach may be advantageous if memory transfer between the GPU and CPU is not the rate-limiting step. Furthermore, in this case the GPU can also optimize computing timing by queueing certain steps for execution while other memory is being transferred. We found that configuring NeuroGPU in SingleKernel mode produced the fastest runtimes in all models tested (Table 4), and had higher GPU utilization levels. This indicates that, for most models, memory transfer between GPU and CPU is rate-limiting, and models run most efficiently when the majority of calculations are isolated on the GPU. Nevertheless, the highest utilization values were ~10% in the SingleKernel configuration (3.8% in SplitKernel), suggesting that additional memory optimizations could be leveraged in future iterations of NeuroGPU.

**Table 4:**

| Model | Morphology | SingleKernel |  | SplitKernel |  |
| --- | --- | --- | --- | --- | --- |
|  |  | Acceleration | Utilization | Acceleration | Utilization |
| Passive membrane | Soma & apical dendrite | 95.8 | 0.30% | 78.8 | 0.41% |
|  | Pyramidal Cell | 58.1 | 10% | 55.2 | 3.62% |
| Mainen and Sejnowski | Soma & apical dendrite | 153.1 | 10% | 99.5 | 1% |
|  | Pyramidal Cell | 114.2 | 10% | 111.8 | 3.80% |
| Blue Brain Project | Pyramidal Cell | 93 | 10% | 107 | 3.70% |
|  | Chandelier Cell | 205.6 | 10% | 197.5 | 1.60% |
| Acceleration: fold increase in processing speed relative to single core CPU (MODEL) |  |  |  |  |  |
| Utilization: Percent of time the GPU is being used |  |  |  |  |  |

## Results

Our goal for NeuroGPU was to develop user-friendly software for fitting compartmental models to experimental data, with improved speed, using relatively low-cost hardware. Furthermore, we sought to make NeuroGPU cross-compatible with NEURON, to allow one to import models available on public databases, including ModelDB and the BBP portal. Toward that end, we utilized the same basic structure as NEURON, including the use of *hoc* and *mod* files that define all aspects of the compartmental model (Fig. S1). To increase simulation speed, we focused primarily on parallelizing the most computationally intensive aspects of NEURON simulations in GPU architecture. NEURON calculates the voltages of each segment of the model by solving a system of differential equations that describes current flow in each compartment. Within NEURON, this system of differential equations is represented within a tri-diagonal matrix (Hines, 1984). Typically, matrix elements for neighboring compartments are solved in serial, as current flow in one compartment is interdependent on flow in neighboring compartments. We and others have previously developed methods to solve this tri-diagonal matrix in parallel across GPUs, despite the interdependence of current flow across compartments (Fig. S2) (Hines, 1984; Hines et al., 2008; Ben-Shalom et al., 2013). At that time, the method was implemented only for classic Hodgkin-Huxley models with 3 parameters (gNa, gK, gLeak) (Ben-Shalom et al., 2013). Here, we extended this method to support a wider range of models, including most models available in ModelDB and the Blue Brain Project (BBP) repository. We implemented this in Python and created an iPython Graphical User Interface (GUI) for easy use.

### NeuroGPU Performance

To evaluate how NeuroGPU performs relative to NEURON, we benchmarked it for speed and accuracy across different conditions, including CUDA implementation, hardware configurations, and across a range of models. We first compared NeuroGPU performance with a single GPU to NEURON implemented on a single CPU core. To benchmark speed, we evaluated computing time for multiple instances of the same model. NeuroGPU was primarily evaluated on recently developed models from the Blue Brain Project (BBP) portal (Hay et al., 2011; Markram et al., 2015; Ramaswamy et al., 2015), but was also benchmarked on models with reduced morphology or reduced numbers of voltage-gated channels or ligand-gated receptors to determine how each of these aspects affects performance (Figs. S3-4). For BBP models, we focused on two specific models: a layer 5 pyramidal neuron (Fig.2, top row: BBP_PC, see Methods for specific model) and a layer 5 chandelier interneuron (Fig.2, bottom row: BBP_CC). Models were interrogated with a range of stimulus intensities to determine relative differences between NeuroGPU and NEURON (Fig. 2).

We first confirmed the quality of the simulations. Overall, NeuroGPU was able to replicate NEURON simulations with high fidelity; however, small voltage differences were observed between the two platforms in all models tested. These were most commonly observed when voltage was changing rapidly between time steps (Fig. 2B-C, G-H), and were due to small differences in timing that likely arise from different approaches to number rounding in GPUs vs CPU architecture (Whitehead, 2011).

NEURON computation time scales linearly with the number of simulations, and, for relatively small numbers of models (< 8), outperforms NeuroGPU (Fig.2, D,E,I,J). By contrast, models implemented on GPUs scale linearly only after saturating all of the GPUs streaming multiprocessors (Nvidia, 2018) (Fig.2, D,E,I,J). With NeuroGPU, processing times are quite similar for any simulation incorporating fewer than 128 models, and begin to outpace NEURON simulations when >8 simulations are run simultaneously. Relative gains in processing time were noted when 8 to 16,384 models were run simultaneously. These gains were dependent on hardware (Fig.2, E,I). For example, implementing NeuroGPU on an NVIDIA TitanXP GPU resulted in 46.6-fold improvements in processing speed, while the same models run on an NVIDIA Tesla V100 were 205.6-fold faster. It is worth noting that TitanXP hardware is relatively low cost (<$1099) and a very similar card (NVIDIA GTX-1080) can currently be purchased for less than $500. As such, significant improvements in processing speed can be obtained even with modestly priced hardware. Additional returns can be gained from GPU tethering, as CUDA has been recently updated to allow for memory sharing across GPUs. To evaluate this, we connected up to 4 Tesla V100 GPUs together and measured speedup on both BBP models displayed in Figure 2. As expected, adding more GPUs increased the overall processing capacity, and we noted shifts in the number of models that could be handled simultaneously before reaching maximum GPU utilization (Fig. 2E, J). Furthermore, speedup was almost 3 orders of magnitude faster relative to NEURON.

### Exploring neuronal model parameter space

Parameter values (e.g. ion channels distributions) are correlated in a non-linear manner. This may lead to situations where vastly different parameter combinations nevertheless produce similar voltage outputs, at least for a limited set of stimuli (Prinz et al., 2004). The diversity of these parameter sets can be limited by constraining the range over which a parameter can vary before initiating model optimization, thus leading to more biologically realistic sets of parameters. NeuroGPU may be ideal for parameter exploration within such ranges, as these types of simulations require one to repeatedly model the same morphology with small differences in constituent parameter values, a process that lends itself well to parallelization within GPUs. Indeed, we predict that relative speedups would be identical to situations considered above (Fig. 2) and depend simply on the number of parameter sets used. To provide an example of parameter space exploration, we examined neuronal output (i.e., number of action potentials) in the BBP_PC model when co-varying the density of the axonal fast inactivating sodium channel and axonal slow-inactivating potassium channel over a range of 0 to 10 and 0 to 20 S/cm^2^, respectively. Single traces from with different sodium and potassium conductances are shown in Figure 3A and total spike output as function of these two channel densities is shown in Figure 3B. As expected, increasing sodium conductance allowed models to generate more APs until sodium conductance was so high that models entered depolarization block. Similarly, reducing potassium conductance produced comparable results. Interestingly, certain combinations of sodium and potassium conductance concentrations produced bursting phenotypes characterized by high-frequency APs riding atop long-duration depolarizations. These presumably reflect parameter ranges that then interact with other ion channels in the model (e.g., Ca_V_3 of HCN channels) that promote such burst dynamics.

### Fitting models to surrogate and empirical data

With the ability to rapidly sample parameter space, NeuroGPU may be ideally suited to accelerate model fitting to data, where a key constraint is the time needed to exhaustively sample possible solutions. To test this, we implemented two different genetic optimization algorithms within NeuroGPU. Initially we integrated the DEAP (Distributed Evolutionary Algorithms in Python) package (Gagn, 2012) (Fig. 4). Genetic algorithm success lies in the balance between exploration of the whole parameter space and the exploitation of specific areas that seem promising. For this, large sample populations are ideal, as this allows for effective and broad parameter space exploration. NeuroGPU is more efficient when many instances are running in parallel, allowing for more effective application of genetic algorithms.

**Figure 4:**
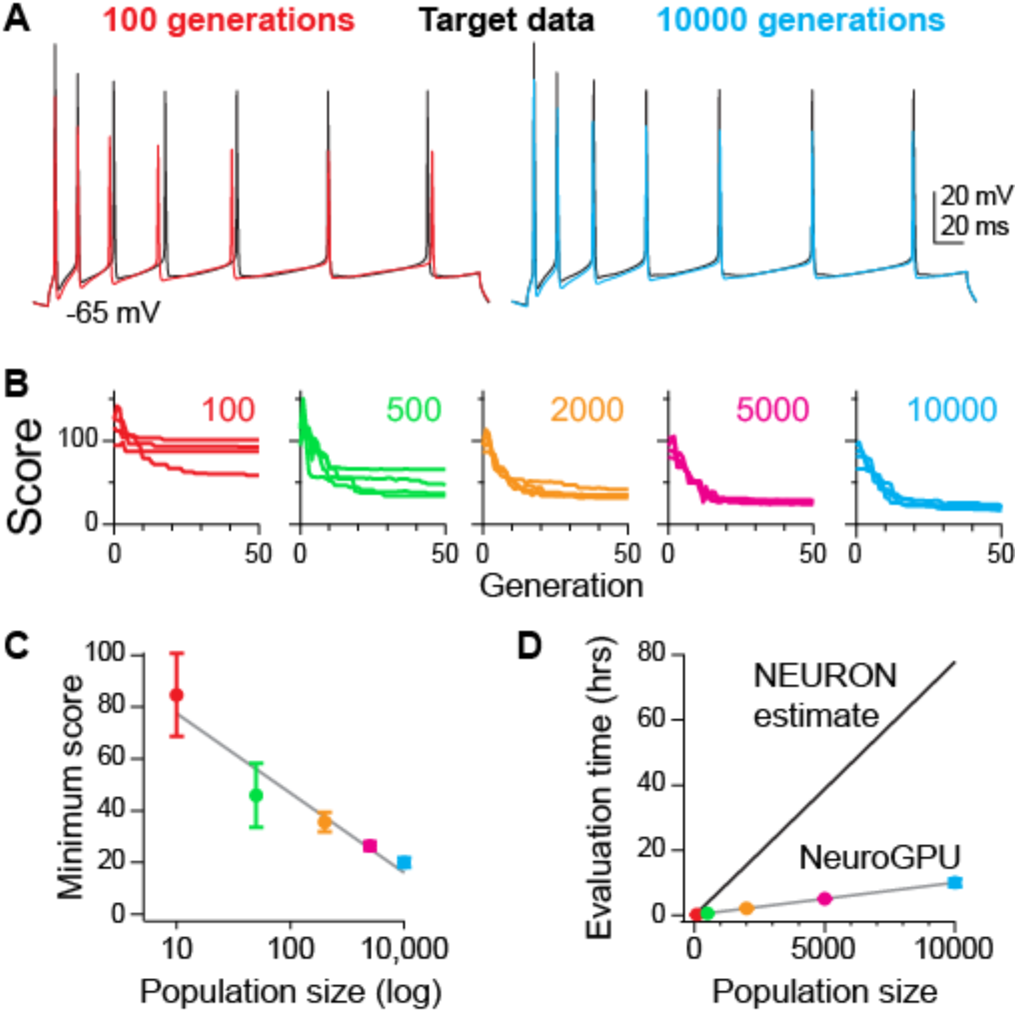
NeuroGPU accelerates evolutionary optimization for fitting models to data. **A:** Voltage traces obtained from optimization (worst case from population of 100: red; best case from population of 10,000: cyan) compared to ground truth (black). **B:** Optimizations examples using DEAP with different sizes of populations. Four Optimizations with different random starting population over 50 generations. Y axis is the error from the target voltage as described in the methods section. Lower values denote less error from target data. **C:** Comparing runtimes for optimizations using NeuroGPU and NEURON (linearly extrapolated from 5 generations). Circles are color coded for population size as in A, and represent mean ± SEM. **D:** Best score in each optimization in A. Circles and error bars as in C.

Genetic optimization was tested here by fitting model-generated voltages to a single voltage epoch containing APs that was generated by the default values present in the BBP_PC model. We focused first on such surrogate data, as the ground truth values for all parameters are already known. As such, we can easily compare how well NeuroGPU performs in arriving at similar values. Optimization began with different population sizes comprised of 100 to 10,000 individual parameter sets with random initial values (Fig. 4B). These populations were run in four independent trials, each for 50 generations, and the difference between the naïve model and ground-truth model was compressed to a single score value (see Methods). For these scores, lower values indicate less difference between the two cases.

Scores improved for each of these populations, but the variance across trials and the overall score were markedly affected by the population size, with score decreasing in a near-linear fashion with each doubling of population size (Fig. 4C). These score improvements were paralleled by a decrease in total processing time. For example, optimization with 10,000 individual parameter sets ran 7.7x faster on NeuroGPU than NEURON (Fig. 4D; 10 vs 77 hours, respectively). While these are significant improvements in simulation speed, they are relatively modest compared to those observed in other conditions (Fig. 2), likely since the version of NeuroGPU used here required NEURON to load the simulation and generate parameter values. In the next section we present how eliminating calls to NEURON greatly increases speedup.

Given these promising results fitting surrogate data, we next asked whether similar performance could be noted for empirical electrophysiological data. Therefore, we fitted the BBP PC model to whole-cell current-clamp recordings of action potential activity from neurons in acute mouse frontal cortex slices (Fig. 5A black traces). Models were implemented on two different pyramidal cell morphologies from somatosensory (original BBP PC morphology; Fig. 5A) and prefrontal cortex [from neuroMorpho.org; Fig. 5B (Ascoli et al., 2007; Yin et al., 2018)] to determine whether morphology differentially affects optimization. We fitted the model to eight voltage responses from varying stimulations to verify that the model is robust across stimulation conditions (Fig. 5A, B). The optimization algorithm found values for the model’s parameters that resulted with good fits to both morphologies. Thus, models can be fitted to empirical data and new morphologies by combining NeuroGPU with BluePyOpt.

**Figure 5:**
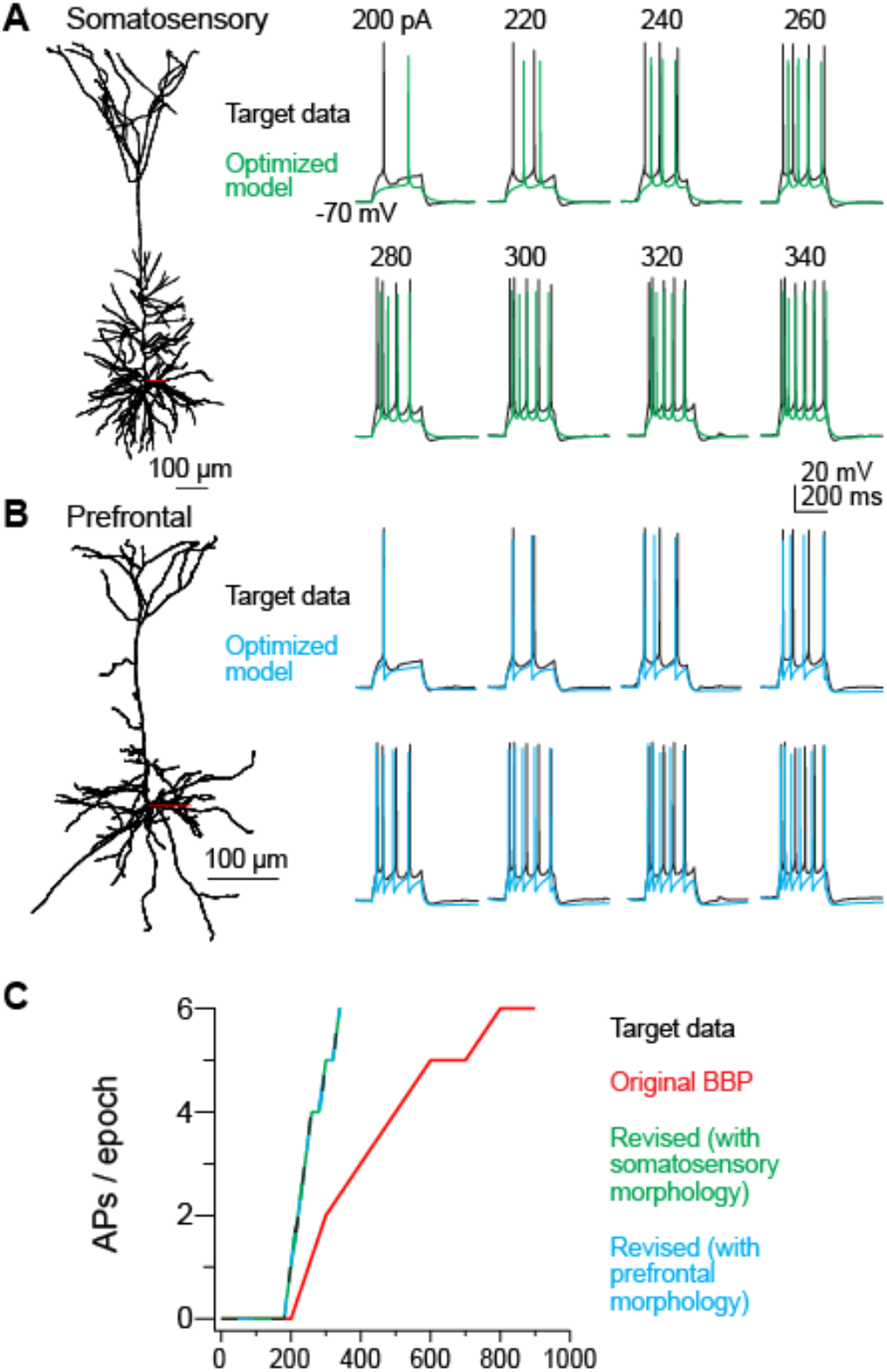
NeuroGPU fits BBP PC model to empirical data. **A:** Left: morphology of L5 thick-tufted pyramidal neuron from somatosensory cortex (Ramaswamy et al., 2015). Right: NeuroGPU fits (green traces) the L5 PC model to empirical data recorded from an L5 prefrontal cortex pyramidal neuron of a mouse (Black Traces) with different stimulus intensity 200-340pA (Spratt et al., 2019) **B:** Left: Prefrontal-cortex layer 5 pyramidal neuron morphology (Ascoli et al., 2007; Yin et al., 2018). The morphology of BBP PC model was modified to that of the prefrontal neuron and fitted to the same data as in A. **C:** Number of APs per 300 ms of step stimulation for different models of layer 5 pyramidal neuron. Note that both revised models overlap target data.

Unlike single trace stimulations (Fig. 4), optimization to 8 separate traces required more generations to obtain reasonable fits, which unfortunately increased processing time dramatically. To reduce runtimes, we revised NeuroGPU by adding two final features. First, we eliminated NEURON calls to the CPU entirely by generating a procedure that identifies the free parameters of the model and modifies NeuroGPU’s input, ‘AllParams.csv’, to new values without using NEURON. Second, we used CPU multithreading scoring, which reduced the overall computation time of each generation. With these improvements, runtimes were reduced for each iteration of the optimization algorithm. Simulation of all 8 traces with 10,000 putative solutions required 60 seconds of processing time. This was followed by a 103 second period required for scoring for each genetic algorithm generation. This latter aspect has been accelerated dramatically. On 8 CPUs, the same process requires 2891 seconds per generation. These speedups (48.2 fold in simulation time) are close to the speedups of single simulations (Fig. 2D). With these improvements, we were able to obtain models that fit well to experimental data using an 8 GPU node running NeuroGPU in just 4 hours, a task that traditionally would require several days of computation using a large cluster (Hay et al., 2011; Hill et al., 2011; Almog and Korngreen, 2014). As such, NeuroGPU allow one the ability to develop more complex models and fit them to empirical data in reasonable time frames (Fig. 1).

## Discussion

Detailed models of neurons are critical to our understanding of neuronal functioning. However, the computational resources required of current software implementations of complex neuronal models are prohibitive, typically requiring supercomputers. At best, this limits the accuracy of results; at worst, it limits access to all but a select set of scientists. To address this gap, based on our previous efforts (Ben-Shalom et al., 2013), we designed a user-friendly environment that enables one to port multi-compartmental models for implementation with CUDA to run simulations on GPUs. By taking advantage of parallel processing inherent to GPUs, we were able to accelerate simulations dramatically—up to 2-3 orders of magnitude with multiple GPUs. NeuroGPU was developed to be interoperable with NEURON, thereby allowing anyone with expertise in the NEURON environment access to GPU-based acceleration. Towards this goal, we developed a platform to easily port NEURON models from either ModelDB or the BBP portal (Ramaswamy et al., 2015; McDougal et al., 2017) using a iPython notebook-based graphical user interface (GUI). We further developed GUIs for creating stimulation protocols, parameter exploration, and genetic optimization.

NeuroGPU addresses a major gap in currently implemented GPU-based simulation environments. Two other neuronal simulations environments for multi-compartmental models have been implemented using GPUs, CoreNeuron (Kumbhar et al., 2019) and Arbor (Akar et al., 2019). These environments are designed primarily to accelerate large scale network simulations. NeuroGPU, by contrast, is focused on greatly accelerating the simulation of single neurons with complex, multi-compartment morphologies, critical for exploring the parameter space of single models and optimizing such models to best fit empirical data.

Leveraging the acceleration of single-neuron simulations, NeuroGPU has expanded GUIs for parameter exploration, which allows for rapid assessment of how changes in ion channel density across compartments affects neuronal excitability (Fig. 3). This approach may be particularly useful to generate testable hypotheses regarding channel distribution with pharmacological manipulations (Keren et al., 2009; Almog and Korngreen, 2014; Mäki-Marttunen et al., 2018), modulation of ion channels (Byczkowicz et al., 2019), or in disease states where ion channel density is thought to be affected (Migliore and Migliore, 2012; Miceli et al., 2013; Ben-Shalom et al., 2017; Spratt et al., 2019). Furthermore, one could generate a range of cells with variable channel densities and confirm that their activity is physiologically realistic (e.g., Fig. 3, all cases before generating depolarization block). These conditions could then be used as building blocks for variable activity within neuronal networks (Prinz et al., 2003, 2004; Alonso and Marder, 2019). In addition to parameter exploration, NeuroGPU is designed for extensive model optimization (Gagn, 2012; Van Geit et al., 2016). Fitting complex models to empirical data is computationally expensive, often requiring days of compute time, even on large supercomputing systems (Hay et al., 2011; Almog and Korngreen, 2014). Here, we show that NeuroGPU accelerates model fitting processing times by 2-3 orders of magnitude (Fig. 4-5). This makes it possible for individual labs to implement optimization algorithms with their own hardware. But also, it opens the door to extremely high-speed model fitting, as NeuroGPU can be run easily on GPU supercomputing systems, which inherently have exponentially more computational resources compared than similarly kitted CPU systems.

Recent advances in genetic characterization and novel analysis methods have resulted in characterization of diverse neuronal types with respect to their morphologies, projections, and protein expression (Gouwens et al., 2019). However, these advancement lack detailed biophysical models that can describe and simulate these neurons. In the BBP portal there are ~200 models that each is comprised of an m-type, which describes the morphology, and one of 11 e-types, which is a set of conductances that describes the electrical properties of the cell. Currently, these e-types describe somewhat generalized activity patterns [e.g., neurons that fire at high frequency without accommodation, or neurons that have stuttering firing patterns (Markram et al., 2015)]. With NeuroGPU we can easily expand this repertoire of e-types, and even replace generalizable e-types with models that recapitulate the activity of data obtained from single neurons. This would improve network simulations, as single neurons would display biologically accurate diversity within and across neuronal subtypes (e.g., Fig. 5B).

NeuroGPU accelerates compartmental modeling through parallelization of matrix calculations. Solving the tridiagonal matrix is the most computationally demanding aspect of multi-compartmental model simulations (Hines, 1984; Hines et al., 2008; Ben-Shalom et al., 2013). Therefore, we took advantage of fast, on-GPU memory and controlled the timing of calculations and memory transfers to optimize the use of computational resources (Volkov and Demmel, 2008; Ben-Shalom et al., 2013; Nvidia, 2018). Resulting speedups depended primarily on neuronal morphology, and in general we show that NeuroGPU offered the best improvements when processing morphologically complex cases. Even in these cases, overall GPU utilization was limited by execution dependencies, where one aspect of GPU processing could not proceed until another aspect either transferred or processed its own memory. In the future, these dependencies may be further reduced through either dynamic parallelization (Zhang et al., 2015) or by increasing instruction level parallelism (ILP) (Volkov and Demmel, 2008). Nevertheless, the current version of NeuroGPU accelerates single neuron compartmental simulations by several orders of magnitude.

Future iterations of NeuroGPU may expand on the strengths and address limitations in using GPUs for compartmental modeling. Ion channels are modeled typically with Markov-based kinetics, or a simpler Markov approximation based on Hodgkin-Huxley type equations. NeuroGPU currently supports Hodgkin-Huxley-based mechanisms only, as we found that implementation of full Markov-based mechanisms on GPUs requires too much shared memory and reduces performance drastically (Ben-Shalom et al., 2012). As with total GPU utilization, improvements in memory handling may improve these cases. Furthermore, GPUs work best when the same instructions are occurring simultaneously on multiple memory addresses. This makes them ideal for iterating through models with identical morphologies and different channel distributions, but less ideal for network models containing a diversity of neuron types. As an intermediate, one could address this limitation by modeling networks containing discrete sets of neurons. For example, a network could contain several compartmental morphology models that each support multiple instances with different channel parameters, similar to the Ring model applied by Arbor (Akar et al., 2019; Kumbhar et al., 2019).

NeuroGPU will help democratize multi-compartmental modeling. While NeuroGPU can support simulations in large, tightly integrated systems using UNIX-based mutli-GPU architectures, it also is ideal for individual laboratories running simulations on Windows-based workstations with GPUs, which are becoming increasingly common. Indeed, a workstation with total costs <$3000, when kitted with appropriate GPUs, can out-perform loosely inter connected CPU-based clusters (i.e., with commodity inter-node connection hardware). This could help broaden the use and utility of multi-compartmental modeling by bringing supercomputer-level processing power to a large range of research settings.

## Acknowledgments

We are grateful to Dr. Gilad Liberman who helped conceptualize this project. To the support and advice of NVIDIA developers – Dr. Jonathan Lefman, Dr. Jonathan Bentz, Dr. Xuemeng Zhang and Angela Chen in optimizing the CUDA code. To Maxwell Chen and Mathew Derango who helped with code development. To NVIDIA Corporation for donating the GPUs used in this study. To all the members of the Bender Lab for critically assessing this work. This research was supported by NIH Grants F32 NS095580 (RBS), MH112729 (KJB), and DA035913 (KJB).

---

**Supplemental Figure 1:**
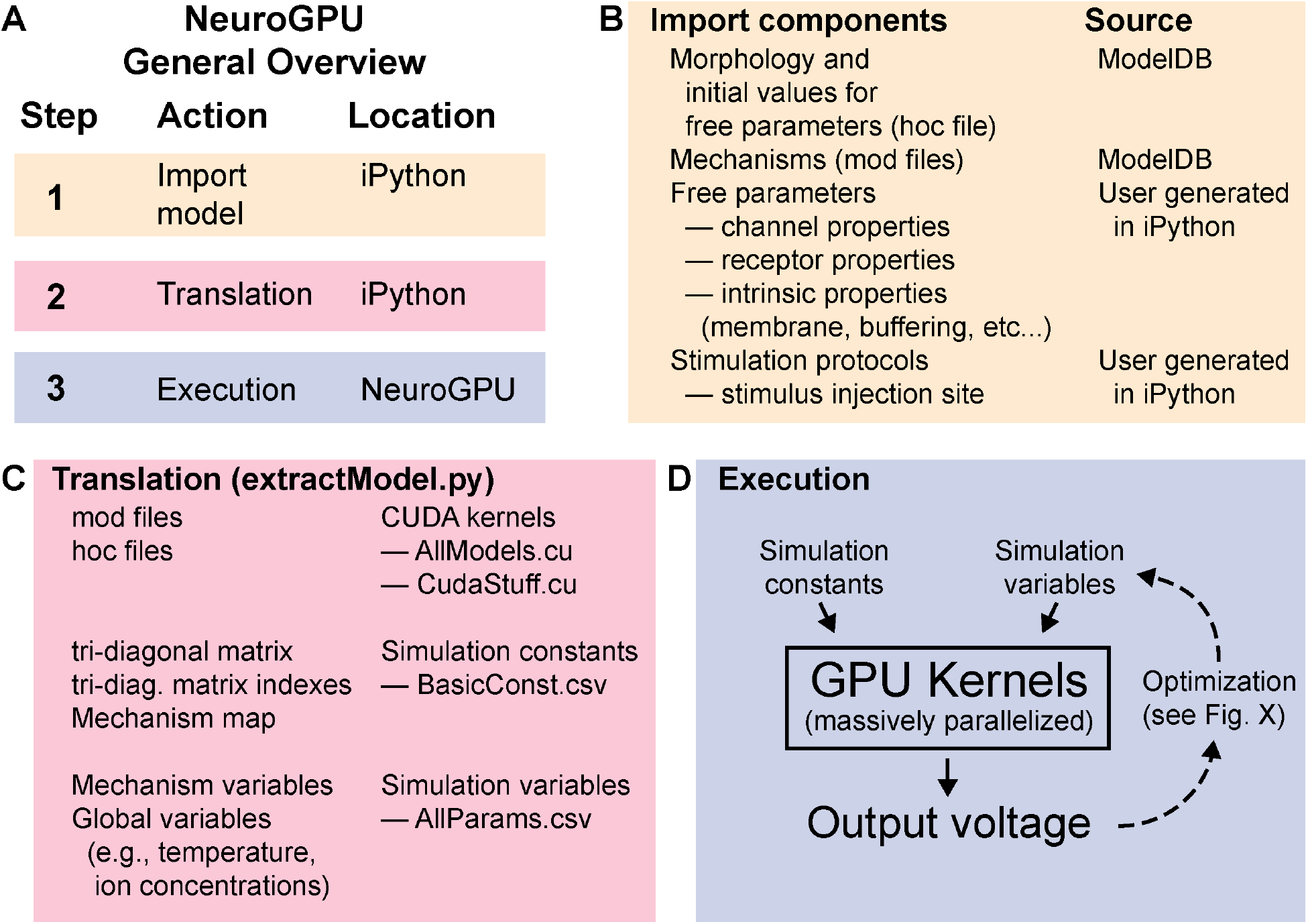
NeuroGPU overview and flowchart. **A:** Overview of the general workflow in NeuroGPU: The user ports a model via the iPython GUI and customizes the simulation (panel B). NeuroGPU translates the model to CUDA code that can run on the GPU and compiles executable code. **B:** Sources for model components: The morphology and model’s properties are described in the hoc file. Additional mechanisms such as ion-channels are described in .mod files. The stimulation protocols can be either imported or can be generated with our provided GUI **C:** Import to NeuroGPU is done by the extractModel.py script. It translates mod files to GPU kernels (see methods), which are written to AllModels.cu, and updates the course of the simulation at CudaStuff.cu. extractModel.py writes to the BasicConst.csv the tri-diagonal matrix and mechanism map, which indicates the mechanisms for each compartment. Finally, extractModel.py writes all the mechanism parameters to AllParams.csv. **D:** After extractModel.py terminates, it creates NeuroGPU.exe. When NeuroGPU is invoked it reads the input files and runs the simulations for the different instances of the model and writes their voltages output to a file. When NeuroGPU is used for optimization, new instances of the models are created each iteration, and only AllParams.csv is updated via a python script.

**Supplemental Figure 2:**
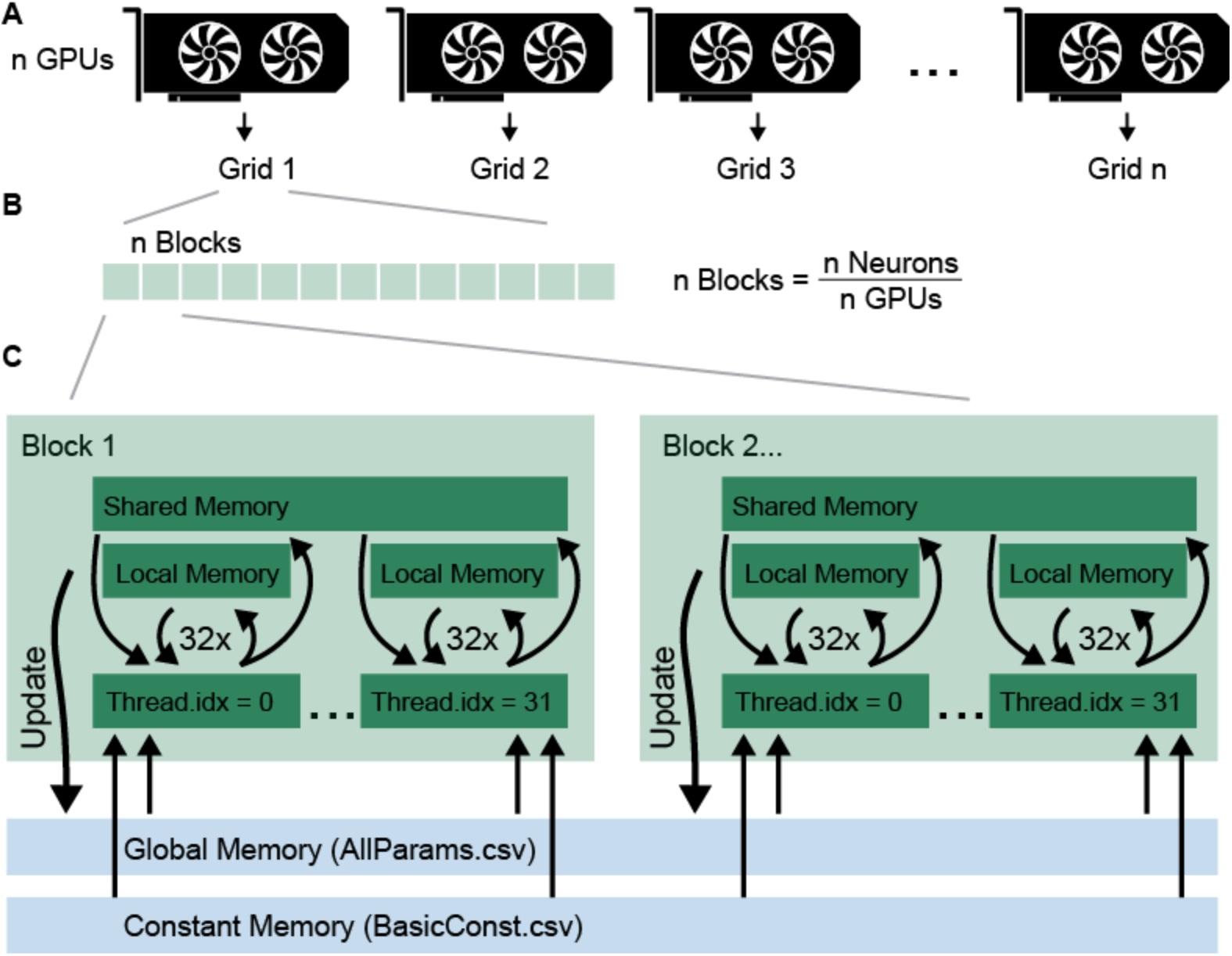
NeuroGPU CUDA implementation. **A:** NeuroGPU can be run on multiple GPUs; each GPU will run a separate grid of block/neurons (Nvidia, 2018). **B:** Grids are distributed in blocks, with each block representing an instance of a model. The number of blocks in a grid is set by the number of model instances that will be simulated on an individual GPU. **C:** A block is the basic simulation unit upon which 32 threads each update the memory in an ILP manner (see Methods). Global memory, which can be accessed by all blocks, stores mechanism parameters for every compartment. Constant memory, which is limited in size, stores the simulation constants such as the tri-diagonal matrix and the mechanism map.

**Supplemental Figure 3:**
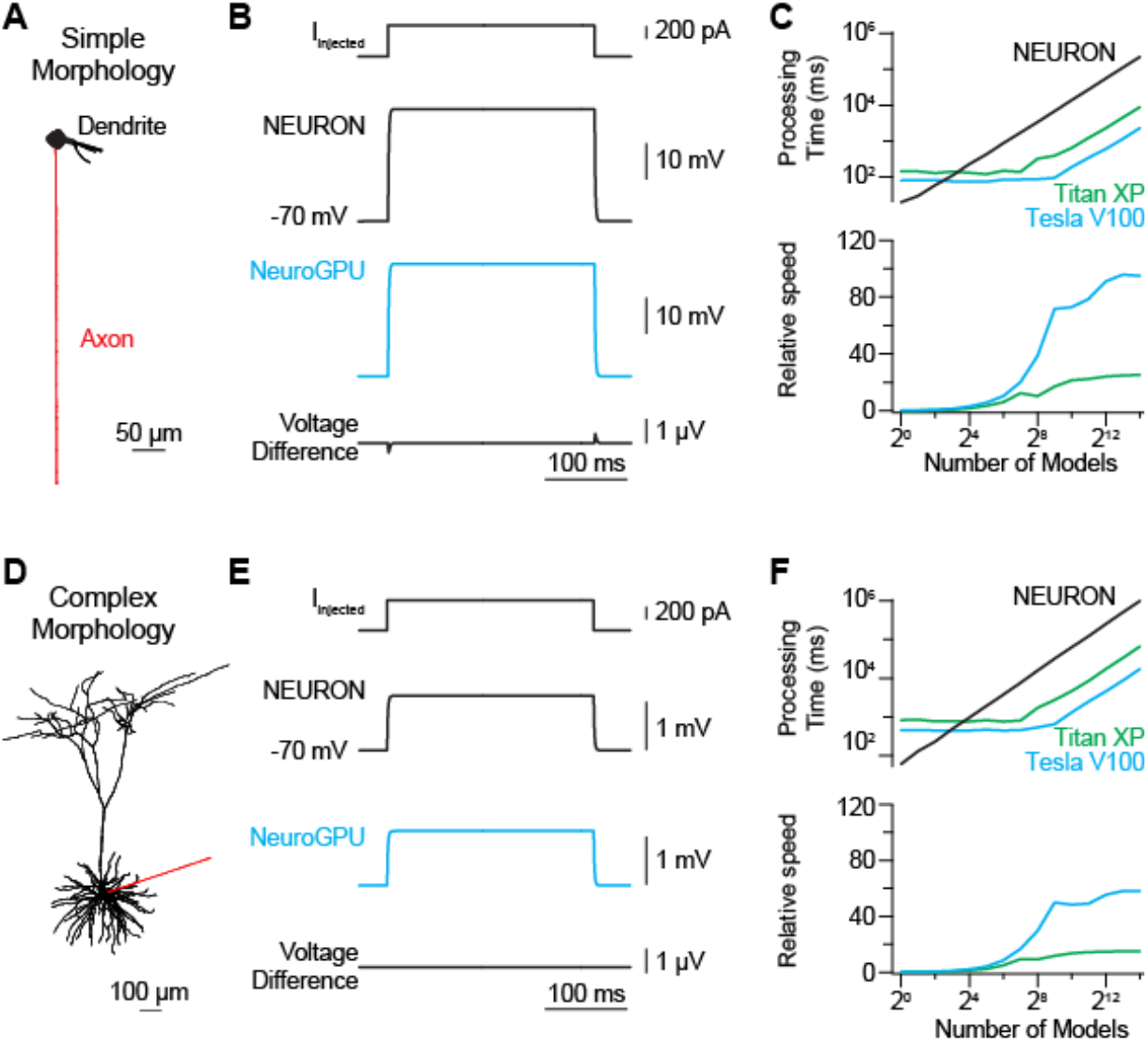
Passive model simulations. **A:** Simple morphology with artificial axon. This model contains passive channels (pas.mod) in all compartments. **B:** Top: injected current at the soma Middle: NEURON voltage response as recorded at the soma. Blue: NeuroGPU response as recorded at the soma. Bottom: difference in voltage between NEURON and NeuroGPU. **C:** Top: Runtimes for the model using the different architectures: black – NEURON, green – NeuroGPU on TitanXP, blue – NeuroGPU on TeslaV100. X-axis in log2 scale, Y-axis in log10 scale. Bottom: Speedup compared to NEURON. **D-F:** Same as A-C, but for complex morphology from (Mainen and Sejnowski, 1996b).

**Supplemental Figure 4:**
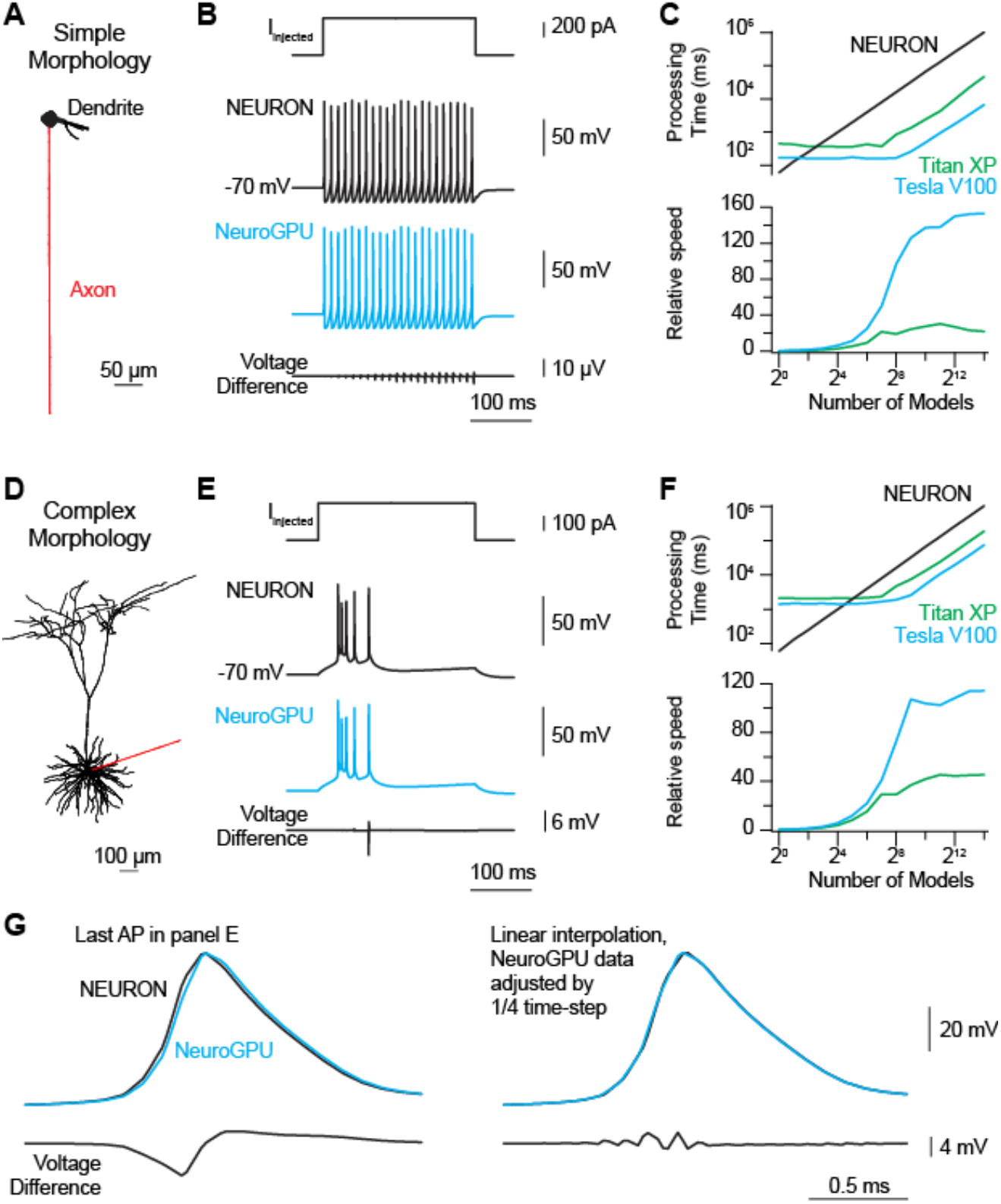
Mainen and Sejnowski model neuron simulations. **A:** Simple morphology with artificial axon and active and passive components distributed as in (Mainen and Sejnowski, 1996b) **B:** Top: injected current at the soma. Middle: NEURON voltage response as recorded at the soma. Cyan: NeuroGPU response as recorded at the soma. Bottom: difference in voltage between NEURON and NeuroGPU. **C:** Top: Runtimes for the model using the different architectures: black – NEURON, green – NeuroGPU on TitanXP, blue – NeuroGPU on TeslaV100. X-axis in log2 scale, Y-axis in log10 scale. Bottom: Speedup compared to NEURON. **D-F:** Same as A-C, but for neocortical layer 5 pyramidal cell morphology, as in (Mainen and Sejnowski, 1996b). **G:** In panel E, a relatively large voltage discrepancy of 6.6 mV was identified. This discrepancy occurred during the last AP within a burst and was due largely to a shift in the timing of this AP (Fig. 4G). Voltage differences are minimized by linearly interpolating the data 4-fold and advancing NeuroGPU simulation by ¼ time-step.

